# Ventral tegmental area dopamine neural activity switches simultaneously with rule representations in the prefrontal cortex and hippocampus

**DOI:** 10.1101/2024.09.09.611811

**Authors:** Mingxin Ding, Porter L. Tomsick, Ryan A. Young, Shantanu P. Jadhav

**Affiliations:** Graduate Program in Neuroscience, Brandeis University, Waltham, MA 02453, USA; Undergraduate Program in Neuroscience, Brandeis University, Waltham, MA 02453, USA; Department of Neuroscience, Virginia Polytechnic Institute and State University, Blacksburg, VA 24061, USA; Department of Psychology, Brandeis University, Waltham, MA, 02453, USA; Volen National Center for Complex Systems, Brandeis University, Waltham, MA, 02453, USA

## Abstract

Multiple brain regions need to coordinate activity to support cognitive flexibility and behavioral adaptation. Neural activity in both the hippocampus (HPC) and prefrontal cortex (PFC) is known to represent spatial context and is sensitive to reward and rule alterations. Midbrain dopamine (DA) activity is key in reward seeking behavior and learning. There is abundant evidence that midbrain DA modulates HPC and PFC activity. However, it remains underexplored how these networks engage dynamically and coordinate temporally when animals must adjust their behavior according to changing reward contingencies. In particular, is there any relationship between DA reward prediction change during rule switching, and rule representation changes in PFC and CA1? We addressed these questions using simultaneous recording of neuronal population activity from the hippocampal area CA1, PFC and ventral tegmental area (VTA) in male TH-Cre rats performing two spatial working memory tasks with frequent rule switches in blocks of trials. CA1 and PFC ensembles showed rule-specific activity both during maze running and at reward locations, with PFC rule coding more consistent across animals compared to CA1. Optogenetically tagged VTA DA neuron firing activity responded to and predicted reward outcome. We found that the correct prediction in DA emerged gradually over trials after rule-switching in coordination with transitions in PFC and CA1 ensemble representations of the current rule after a rule switch, followed by behavioral adaptation to the correct rule sequence. Therefore, our study demonstrates a crucial temporal coordination between the rule representation in PFC/CA1, the dopamine reward signal and behavioral strategy.

**Significance Statement:** This study examines neural activity in mammalian brain networks that support the ability to respond flexibly to changing contexts. We use a rule-switching spatial task to examine whether the key reward-responsive and predictive dopamine (DA) activity changes in coordination with changes in rule representations in key cognitive regions, the prefrontal cortex (PFC) and hippocampus. We first established distinct rule representations in PFC and hippocampus, and predictive coding of reward outcomes by DA neuronal activity. We show that the rule-specific DA reward prediction after a rule switch develops in temporal coordination with changes in rule representations in PFC, eventually leading to behavioral changes. These results thus provide an integrated understanding of reward prediction, cognitive representations of rules and behavioral adaptation.

## Introduction

Behavioral flexibility is critical to survival and for adapting to a changing environment. These functions are frequently driven by changing reward conditions and supported by distributed network processing in the brain to evaluate context and respond appropriately. The hippocampal cognitive map represents spatial context (O’Keefe and Dostrovsky, 1971; O’Keefe, 1978), and hippocampal neuronal activity can also be modulated by rewards and goals (Lee et al., 2012; Gauthier and Tank, 2018; Krishnan et al., 2022). The prefrontal cortex (PFC) serves complementary cognitive functions including working memory, executive control, and context representation (Ragozzino and Kesner, 1998; Yoon et al., 2008; Horst and Laubach, 2009; Durstewitz et al., 2010; Hyman et al., 2012; Urban et al., 2014; Ma et al., 2016). The hippocampus and prefrontal cortex have been demonstrated to coordinate temporally for adaptation of behavioral strategy when experiencing environmental/contextual changes (Guise and Shapiro, 2017; Hasz and Redish, 2020). However, much remains unexplored about how reward signals, acting as major feedback to animals’ state/action choices, coordinate with this navigation and memory system for cognitive flexibility and behavioral adaptation. The dopamine (DA) signal from the midbrain is known to be involved in reward prediction error processing (Schultz et al., 1993, 1997) and value estimates (Roesch et al., 2007; Howe et al., 2013; Hamid et al., 2015; Dabney et al., 2020), and is causally linked to learning (Steinberg et al., 2013; Hamid et al., 2015). DA release profile and firing activity has also been reported to ramp up during reward approach and scale positively with reward quantity and probability (Howe et al., 2013; Engelhard et al., 2019; Krausz et al., 2023). How this DA signal changes during rule switches to support behavioral flexibility at short time scales of a few tens of trials over which behavioral change is observed remains less explored.

In this study, we were interested in examining the dynamics of rule and context representations in the hippocampal (area CA1) and prefrontal regions during reward-guided behavioral adaptation in response to rule switching, and whether these dynamics are related to ventral tegmental area (VTA) DA neuronal activity. The primary hypotheses we aimed to test are whether changes in reward-associated DA neuronal firing activity are temporally coordinated with changes in rule representations in these cognitive regions as animals adapt to changes in contingencies over a few trials due to rule switching, thus supporting a role in cognitive flexibility. The PFC receives input from and sends output to midbrain DA neurons (Carr and Sesack, 2000; Beier et al., 2015; Kabanova et al., 2015; Morales and Margolis, 2017), and PFC and DA are known to contribute to reinforcement learning in a complimentary manner, with prefrontal signals encoding predictive values and dopamine encoding prediction errors (Lak et al., 2020). In addition, DA projections to dorsal hippocampus have been reported (Gasbarri et al., 1994), and CA1 place fields have been shown to change stability based on reward conditions and DA input (Martig and Mizumori, 2011; McNamara et al., 2014; Krishnan et al., 2022). It remains unknown if and how DA signal changes coordinate with rule and context representation in the CA1 and/or PFC, and the time scale of such coordination if it exists.

To investigate these questions, we implemented a novel rule-switching spatial task for rats in a W/M maze and recorded activity simultaneously from CA1, PFC and VTA ensembles as animals performed this task. We found single cell and ensemble codes of underlying rules in both the CA1 and PFC regions. VTA reward signaling and predictive coding of reward outcomes during maze running for the two rules were confirmed. We found that this rule-specific predictive feature of DA spiking activity develops together with correct PFC and CA1 rule decoding a few trials after a rule switch, followed by a change of behavioral strategy within a few trials. Together, this work establishes that the dynamics of DA spiking activity and PFC and CA1 rule representation changes occur in a coordinated manner during a rule switching task.

## Materials and Methods

### Animals and experimental design

Four adult male TH-Cre rats (450-600 g, 3-7 months, RRID: RRRC_00659) (Witten et al., 2011) were used in the current study for behavior and physiology data. All procedures were conducted under the guidelines of the US National Institutes of Health and approved by the Institutional Animal Care and Use Committee at Brandeis University. Animals were bred in house, kept under a 12h/12h light/dark schedule, with ad libitum food and water access till at least 10 weeks old prior to behavioral training and experiments. After daily handling and habituation to a sleep box (30cm long, 30 cm wide, 50 cm tall), animals were moderately food deprived to 85-90% of their initial weight to motivate reward-seeking behavior on an elevated linear track. Once reaching the criterion of obtaining 60 condensed milk rewards in a 20-min session, animals were allowed free access to food for at least a week before surgery. During surgery, virus AAV5-EF1a-DIO-hChR2(E123T/T159C)-mCherry was injected into VTA and a custom multi-tetrode microdrive with optrodes was implanted into CA1, PFC and VTA (see **Surgical procedures**). Tetrodes and optrodes were gradually lowered into the target regions in the time span of 3-4 weeks to allow virus expression while animals recovered and retrained on the linear track. Recordings were performed during the process of learning and rule switching on the W track (see **Behavioral paradigm**).

### Behavioral paradigm

Post-recovery animals learned to perform a rule switching task on the W track (80 cm long of each arm, 7 cm wide, see **Figure 1A**). The task consists of two rules with similar structure but different sequence assignments. The first rule is a standard W track alternation task: animals must travel to the center arm (home) to obtain a milk reward when they set out from a side arm (inbound trajectories), whereas they need to alternate between the left and right arms when they start from the center (outbound trajectories) (Jadhav et al., 2012, 2016; Maharjan et al., 2018; Shin et al., 2019). The second rule presents altered identities between the left and center arm. Therefore, the left arm becomes the new home, and animals must alternate between the center and right arms for outbound trajectories. Once animals learned both rules, they were subject to rule switching in blocks if performance of the current rule reached 80% correct. Rule switching happened without external cues, requiring the animals to use only reward feedback to deduce the change and switch to the optimal strategy for the new rule. During a recording day, animals ran 3-4 epochs of 20-25 min duration. The run epochs were interleaved by 20-40 min sleep epochs. It typically took animals 6-12 running epochs to learn each rule, and an additional training of at least 3-4 days before they could switch rapidly between rules (3-4 switches/rule blocks per day).

**Fig 1:**
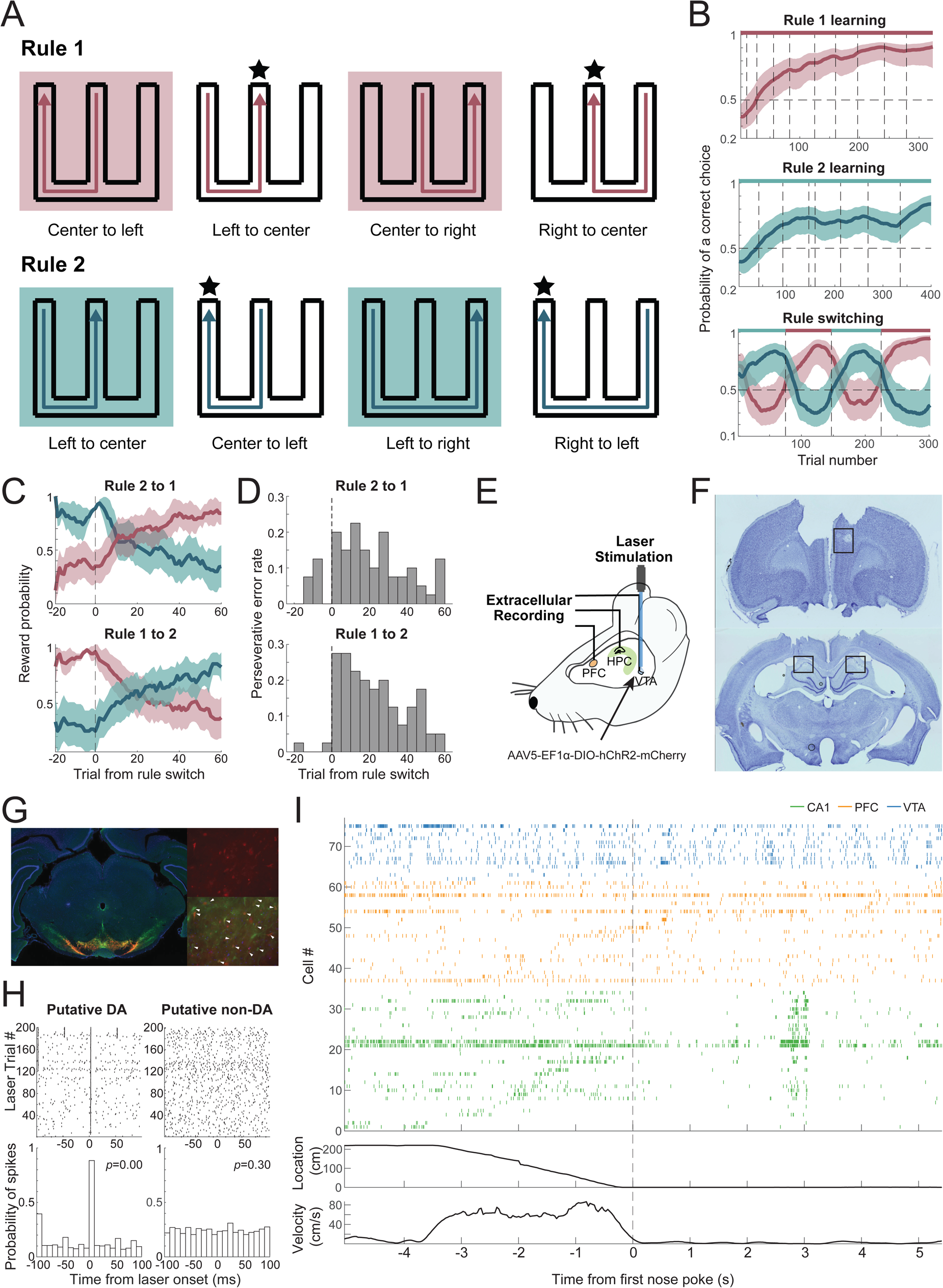
Behavior and Recording Paradigm. A. W-track rule-switching behavior. Top: sequence of rule 1, left-center-right-center; bottom: sequence of rule 2, center-left-right-left. Stars denote the home arm in the given rule sequence. Trajectories in shaded booxes are outbound and require working memory. The common trajectories across the two rules are Left-to-Center and Center-to-Left, with different memory demands for the rules. B. Example performance curve of one animal (estimated mean ± 95% CI). *Left*: Rule 1 learning across behavior epochs/ sessions; *Middle*: Rule 2 learning; *Right*: rapid switching between the two rules within a single day. Note that the rule switch is not indicated by any external cue. The bar on top indicates the current rule (dark red for rule 1, teal for rule 2). Dashed lines in all three panels indicate epoch changes. C. Average reward rate of behavioral choices based on each rule’s reward contingency, for Rule 2 to 1 and Rule 1 to 2 switches separately (n=8 each), aligned to rule switch trials. D. Probability of perseverative error aligned to rule switch trials, for Rule 2 to 1 and Rule 1 to 2 respectively. E. Recording setup. Simultaneous recording in dCA1, mPFC and VTA regions during rule switching behavior, with photo-tagging of TH+ neurons in the VTA in TH-Cre animals. F. Example of Nissl-stained PFC and CA1 slices. Rectangles indicate regions of lesion marks. G. Histology in VTA. Red: virus expression of AAV5-EF1a-DIO-hChR2(E123T/T159C)-mCherry; green: antibody staining of TH+ cells; blue: DAPI. White triangles show overlap between virus expression and antibody staining. H. VTA spiking responses to laser stimulation at 0 ms. Top: raster plot showing individual spikes aligned to each stimulation onset. Bottom: probability of spiking. Left: example of an opto-tagged neuron (p=0); right: example non-tagged VTA neuron (p=0.30, Stimulus-Associated spike Latency Test, SALT). I. Top: spike raster during an example trial starting with the animal leaving the last reward location, running a trajectory to the next reward, followed by immobility and consumption at new reward location. Green: CA1 cells; orange: PFC; blue: VTA. Middle: linear distance to reference reward location. Bottom: movement speed. Dashed line: time of first nose poke at the destination reward location.

**Fig 2:**
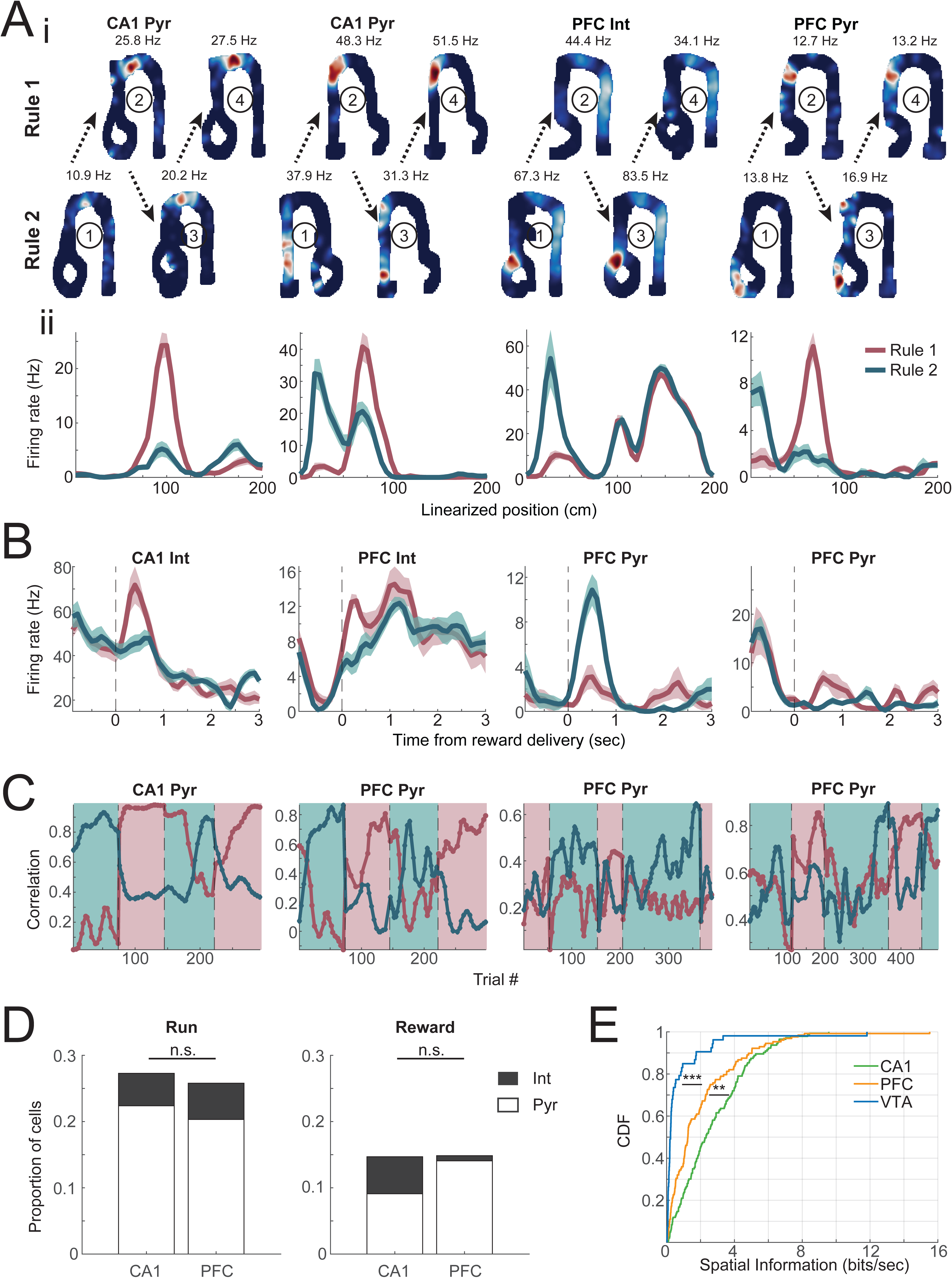
Remapping of single cells in CA1 and PFC across rules. A. Illustrative examples of spatial representations in CA1 and PFC showing remapping on the common trajectories performed across the two rules, Left-to-Center or Center-to-Left. (i). 2D place field at stable performance of rule 1 (*top*) and rule 2 (*bottom*) in interleaved blocks (first block for each rule showed on Left and second block shown on Right, numbers in circles denote order of blocks), with peak firing rate on top. Numbers in circles denote sequence of rule block; (ii). linearized firing fields of the cells shown above (mean ± standard error). B. Illustrative examples showing firing rate change of neurons at the same well locations across the two rules, aligned to reward onset for the same common trajectory across rules (Left-to-Center or Center-to-Left). C. Correlation of single cell firing patterns during individual trials to average firing patterns during stable performance of a given rule (> 60^th^ percentile of the performance of the day) during rule switching blocks. Note the change in correlations that occur after rule switch. D. Proportion of remapped cells for each region. During running: CA1 27.3% (39/143, 32 Pyr, 7 Int), PFC 25.8% (33/128, 26 Pyr, 7 Int), χ^2^=0.077, p=0.78; At reward location: CA1 14.7% (21/143, 13 Pyr, 8 Int), PFC 14.8% (19/128, 18 Pyr, 1 Int), χ^2^=0.0013, p=0.97, Fisher’s exact test for proportions of remapped cells between regions. E. Distribution of spatial information encoded by individual cells for each brain region. CA1: median 2.3 bits/sec, n=143; PFC: median 1.3 bits/sec, n=128; VTA: median 0.23 bits/sec, n=53. CA1-PFC p=0.0010, PFC-VTA p=6.2×10^−8^, CA1-VTA p=7.7×10^−17^, Kruskal-Wallis test with multi-comparison correction.

### Surgical procedures

Each rat received virus injection and microdrive implantation during the surgery. Microdrive fabrication and implantation procedures were similar to previous reports (Jadhav et al., 2012, 2016; Tang et al., 2017; Shin et al., 2019). Anesthesia was induced by ketamine, xylazine and atropine cocktail and maintained by 0.5% - 2% isoflurane. 500 nL of AAV5-EF1a-DIO-hChR2(E123T/T159C)-mCherry was injected into VTA bilaterally (AP: −5.6 mm, ML: ±1.0 mm, DV: −7.8mm). 24 tetrodes each targeting dorsal CA1 (AP: −3.6 to −4.0 mm, ML: ±2.2 mm, DV: - 2.5 mm) and PFC (AP: +3.0 mm, ML: ±0.9 mm, DV: −2.5 to −3.0 mm), and two optrodes targeting VTA, with each optical fiber surrounded by 8 tetrodes, were encased in a microdrive, and implanted against the surface (for CA1 and PFC) or 2 mm into the brain (for VTA). Animals received post-operative analgesia and were monitored closely for at least a week before food restriction and behavioral experiments.

### Data acquisition and processing

Upon approaching target regions, CA1 was identified by the characteristic EEG features, including sharp-wave ripples (SWRs) and theta modulation. PFC depth was targeted to anterior cingulate cortex (ACC) and prelimbic (PrL) regions. VTA was identified by finding optogenetically tagged cells (see **Cell type identification** below). Electrophysiological data was recorded at 30 kHz using Trodes through a 256-channel headstage (SpikeGadgets, San Francisco, CA). Digital input/output signals of nose pokes and reward delivery were simultaneously recorded. Video recordings of animal behavior were captured at 30 fps with sync’d timestamps to the neural recording. All tetrodes were grounded to a screw above cerebellum. Tetrodes of each brain region were referenced using the group average. Spiking data was bandpass filtered between 600 Hz and 6 kHz, and local field potential (LFP) data was bandpass filtered between 0.5 Hz and 400 Hz and down sampled to 1.5 kHz. Animal’s position was tracked using the cameraModule (SpikeGadgets) to identify the red/green LEDs attached to the headstage and verified by the researcher.

Spikes were clustered using MountainSort4 (Chung et al., 2017) and followed by manual inspection and curation in MountainView. Clusters with isolation score >0.9, noise overlap <0.05 and peak signal-to-noise ratio > 2 were accepted and included in this study.

### Histology

After recording, animals were put under anesthesia and recording sites were lesioned by passing a current of 30 µA through the electrode tips. Animals were then perfused 1-2 days later using 4% formaldehyde and the brains were kept in 4% formaldehyde and 30% sucrose solution until being sliced into 50-µm sections. The VTA slices were immuno-stained for TH and imaged to verify colocalization of virus expression and anti-TH antibody at optrode implantation sites. The CA1 and PFC slices were stained with cresyl violet and imaged to verify tetrode locations.

### Cell type identification

Putative DA neurons were identified using optogenetic tagging at least 3 weeks after surgery to allow virus expression. In the first or last sleep epoch of each recording day, a 473-nm light was delivered to the VTA as 5- or 10-ms pulse trains at 1, 4, 10, 20 and 40 Hz to identify TH+ cells expressing ChR2, similar to previous studies (Cohen et al., 2012; Mohebi et al., 2019; Kim et al., 2020). The power of the laser was calibrated to lie within the range of 5-20 mW/mm^2^ at the tip of the fiber to avoid spike waveform distortion. The light-evoked spike latency was then tested using Stimulus-Associated spike Latency Test (SALT, Kvitsiani et al., 2013). Units with p value <0.001 were classified as putative DA neurons.

CA1 and PFC cells were separated into putative pyramidal neurons and interneurons using k-means clustering with parameters including spike width, peak asymmetry and mean firing rate (Barthó et al., 2004; Sirota et al., 2008; Shin and Jadhav, 2024).

### Data analysis

#### Linearization and normalization

Each trial was defined as the time starting from leaving the last reward well, running on the track and arriving at the current reward well, until the end of the stay at the current reward well. There are six trajectories in total defined across the two rules. Occupancy of positions of each trajectory was binned into 40 equally sized spatial bins (5-6 cm/bin) on the track during running, and then binned for 100-ms temporal bins at the reward wells. The occupancy was then smoothed by a 5-bin wide Gaussian kernel. The firing activity of each cell is binned and smoothed in the same fashion and firing rates are normalized by occupancy. For dimension reduction and population activity analyses, firing rates were z-scored for each cell across all task epochs.

#### Behavior analysis

Behavioral performance of each animal on each recording day was estimated using a previously described state space model (Smith et al., 2004; Kim and Frank, 2009). Average reward rate was summarized for rule 1 to 2 and rule 2 to 1, respectively. For analysis separating performance stages, quantile ranges were set individually for each recording day to have equal number of trials for each performance category (**Figure 3F, 4F**). Behavior strategy was estimated by generating the trajectory transition matrix in a 20-trial block with a 1-trial sliding window. The Pearson’s correlation between each transition matrix and the optimal-strategy transition matrix of each rule was computed to determine which rule the animals are currently following (**Figure 3D, 5D**).

**Fig 3:**
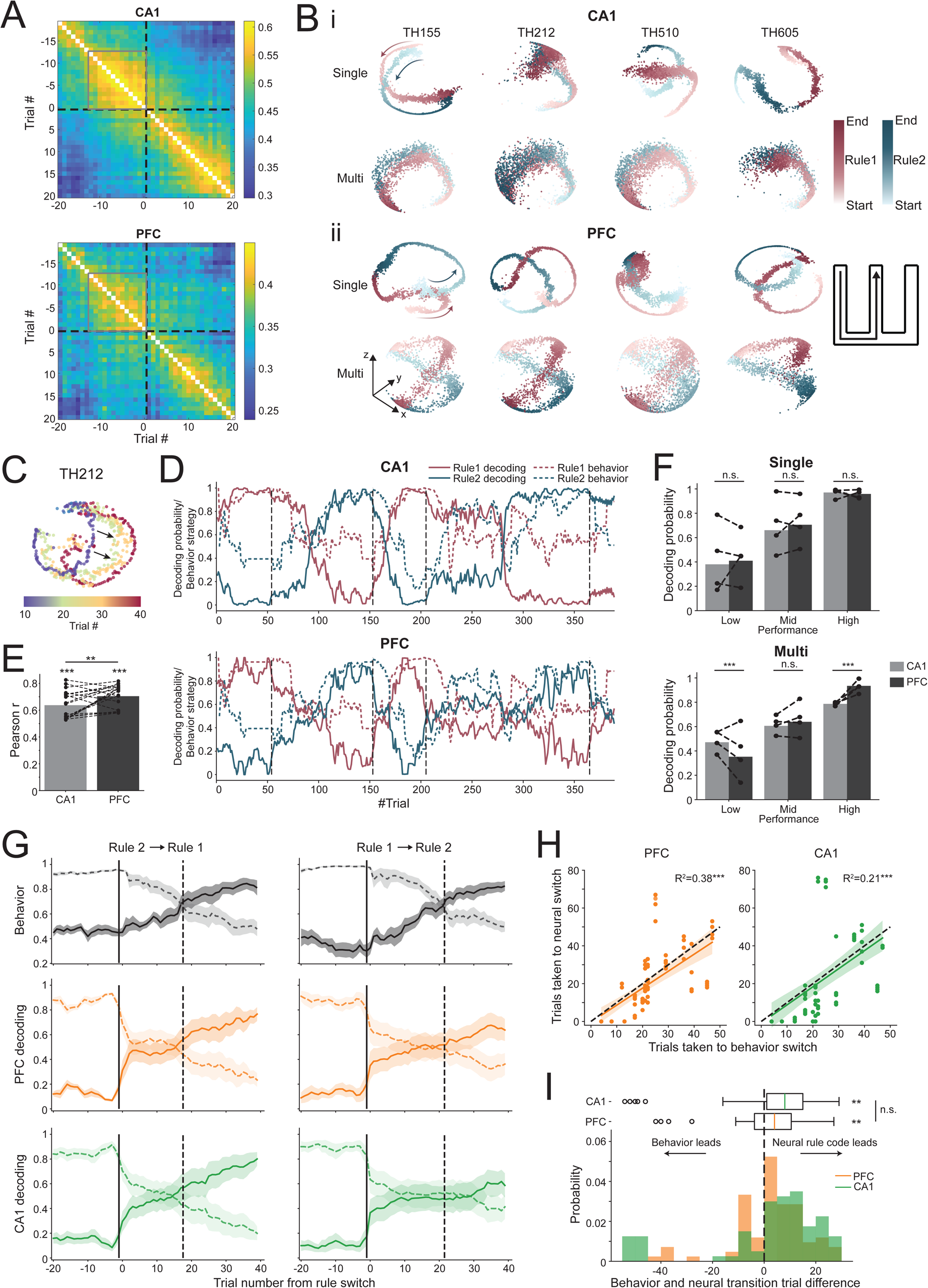
PFC and CA1 ensembles encode rule representations and transitions. A. Correlation of population activity vectors aligned to rule switch (dashed black line) in a trial wise manner for CA1 and PFC. Note the weakened stability shown by lower correlations following rule switches in both regions. Areas of stable correlations are denoted by grey squares. B. CA1 and PFC population activity show differential representation of the same trajectory across rules. Populations activity space shown by dimensionality reduction using CEBRA (see **Methods**). A particular 3D view is illustrated, with arbitrary units for axes. (i) CA1 embeddings of the example common trajectory across rules (Left to Center, schematic on bottom right). *Top*: 3D embedding from models trained using individual animal’s data; bottom: model trained with multi-session CEBRA with data from multiple animals to capture common underlying structure across animals. Start and End denote beginning and end of the same trajectory across the two rules. (ii) Embeddings of the same trajectory but for PFC. C. An illustrative example showing that the PFC population representation of the common trajectory shifts in the manifold space (shift denoted by arrow) during rule transition over the course of trials. Trajectory representations are grouped in 10 trials per color. D. Decoding probability of each rule from an example animal shows CA1 (*top*) and PFC (*bottom*) neural activity transitions from representing one rule to the other (Rule 1 in dark red, Rule 2 in teal) over a few to tens of trials after each rule switching (rule switching is denoted by dashed black vertical lines). Dashed colored curves show estimated behavior strategy similarity to the optimized behavior for each rule. E. CA1 and PFC rule decoding probabilities are highly correlated with behavior strategy. Mean Pearson r: between CA1 and behavior 0.64±0.02, between PFC and behavior 0.70±0.02. Dots and dashed lines show correlation between behavior and each decoding estimate (n=20, 4 animals, 5 estimates each, see **Methods**). p values for all behavior-rule decoding correlations are less than 0.001. Behavior correlation with PFC rule decoding probabilities was higher than with CA1 decoding (p=0.0047, paired t test). F. Rule decoding accuracy (proportion of trials decoded correctly) in each region, grouped by behavioral performance. *Top*: decoding accuracy of single-animal models (low performance: CA1, 0.38, PFC, 0.41, p=0.33 for CA1 vs PFC; mid performance: CA1, 0.66, PFC, 0.71, p=0.12; high performance: CA1, 0.97, PFC, 0.96, p=0.31, two-proportion z-test). *Bottom*: decoding accuracy using multi-animal models (low performance: CA1, 0.47, PFC, 0.35, p=1.3×10^−4^; mid performance: CA1, 0.61, PFC, 0.64, p=0.27; high performance: CA1, 0.79, PFC, 0.93, p=9.9×10^−12^). Dots and dashed lines show decoding accuracy for each animal under each condition (n=4 animals). G. Behavior strategy similarity (top), probabilities of PFC rule decoding (middle) and CA1 rule decoding (bottom) aligned to rule switching trials, shown separately as Rule 2 to Rule 1 (left) and Rule 1 to Rule 2 (right) switches. On average, behavior transitions happened 17 (IQR: 13-29) trials after the Rule1 -> Rule2 switch, with PFC and CA1 neural representations slightly leading behavior. For the switch type of Rule 2 -> Rule 1, behavior started to reflect the new rule after 22 (IQR: 20.75-28.5) trials, with PFC leading in contrast to CA1 lagging. H. Correlation between the number of trials taken for behavior switch and neural representation switch, combined across rule switch types. Dashed lines indicate diagonal lines where the timing of neural transitions equals that of behavior switches. Left: PFC and behavior, R^2^=0.38, p=3.4×10^−10^; right: CA1 and behavior, R^2^=0.21, p=1.9×10^−5^. I. Distribution of timing differences between behavior switch and neural representation switch. Behavior relative to PFC transitions: median +4.0 (IQR: −3.75-10.5) trials, n=80, p=0.0044; behavior relative to CA1 transitions: median +8.0 (IQR: 1-15) trials, n=78, p=0.0040, Wilcoxon signed rank test. For boox plots, booxes, whiskers and circles indicate quartile, 1.5×IQR and outliers, respectively.

#### Spatial information

Spatial information was calculated according to Skaggs et al., 1992 to estimate the amount of spatial content of each cell’s spiking activity:

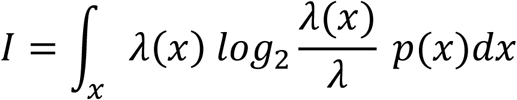

where *I* is the spatial information measure in bits/second, *x* is spatial location of the animal, *p(x)* is the probability of the animal residing at location *x*, *λ(x)* is the mean firing rate of the cell at location *x*, and *λ* is the occupancy-weighted overall mean firing rate of the cell.

#### Dimensionality reduction with Consistent EmBeddings of high-dimensional Recordings using Auxiliary variables (CEBRA)

We used a newly developed nonlinear dimensionality reduction method, CEBRA, to uncover consistent and interpretable latent space of neural spiking data conditioned by behavioral variables (Schneider et al., 2023). Rewarded trials with performance better than 60^th^ percentile of the day were selected for training in CEBRA (**Figure 3B, C**). 20% of these trials with good performance and all trials with worse performance were left out for testing. The training trials were balanced for trajectory identities and rules. We used supervised CEBRA-Behavior models for both single- and multi-animal training. Training labels included spatial-temporal bin, trajectory and rule. Spiking data from each brain region was trained separately for individual animals. Parameters for CEBRA training were largely coherent with the example implementation using rat hippocampus data in (Schneider et al., 2023), except for higher batch size and larger number of iterations to improve model performance:

model_architecture=’offset10-model’,
time_offsets=10,
batch_size=2048,
learning_rate=5e-4,
temperature=1,
output_dimension=3,
max_iterations=12000,
distance=’cosine’,
conditional=’time_delta’,
num_hidden_units=64

The whole data set was then transformed into 3D embeddings using the trained models. To reveal common features across animals, we took advantage of the multi-animal training from CEBRA. This method allows different dimensions (numbers of cells recorded) from different sessions/animals as inputs and maintains the label space across models. Therefore, the resulting embeddings across animals are directly comparable. For multi-animal training, trial selection, model parameters and subsequent transforming process were held the same as in single-session model training.

#### Rule decoding

A k-nearest neighbor (kNN) decoder was trained using the 3D embeddings of training data and used to decode the underlying rule for the testing data. Decoding performance of each trial was calculated as the ratio of bins with correct rule assignment (**Figure 3D**). The decoding accuracy curve was then smoothed using a Gaussian kernel. After a rule switch, a change point was identified as the trial from which decoding probability of the current rule was consistently higher than 50%. This process is done for embeddings from each brain region. We categorized the behavioral performance into low, mid and high by splitting the estimated performance into 1/3 of the trials for each day, and reported the proportion of trials with correct rule decoding in **Figure 3F**. To better estimate the timing of rule representation transitions, we used 5-fold validation for training CEBRA models and decoding, resulting in 5 estimates for each animal and each brain region, reflected in **Figure 3E, H, I**.

#### DA neuron firing rate analyses

Reward responsiveness is defined as showing differences (p<0.05) of firing rates at 200ms – 1200ms after the first nose poke between rewarded and unrewarded trials using Wilcoxon rank sum test. For firing rate comparisons across conditions in **Figure 4C**, firing rates are compared using Wilcoxon signed rank test for each spatial/temporal bin and Bonferroni corrected for multi-comparison.

**Fig 4:**
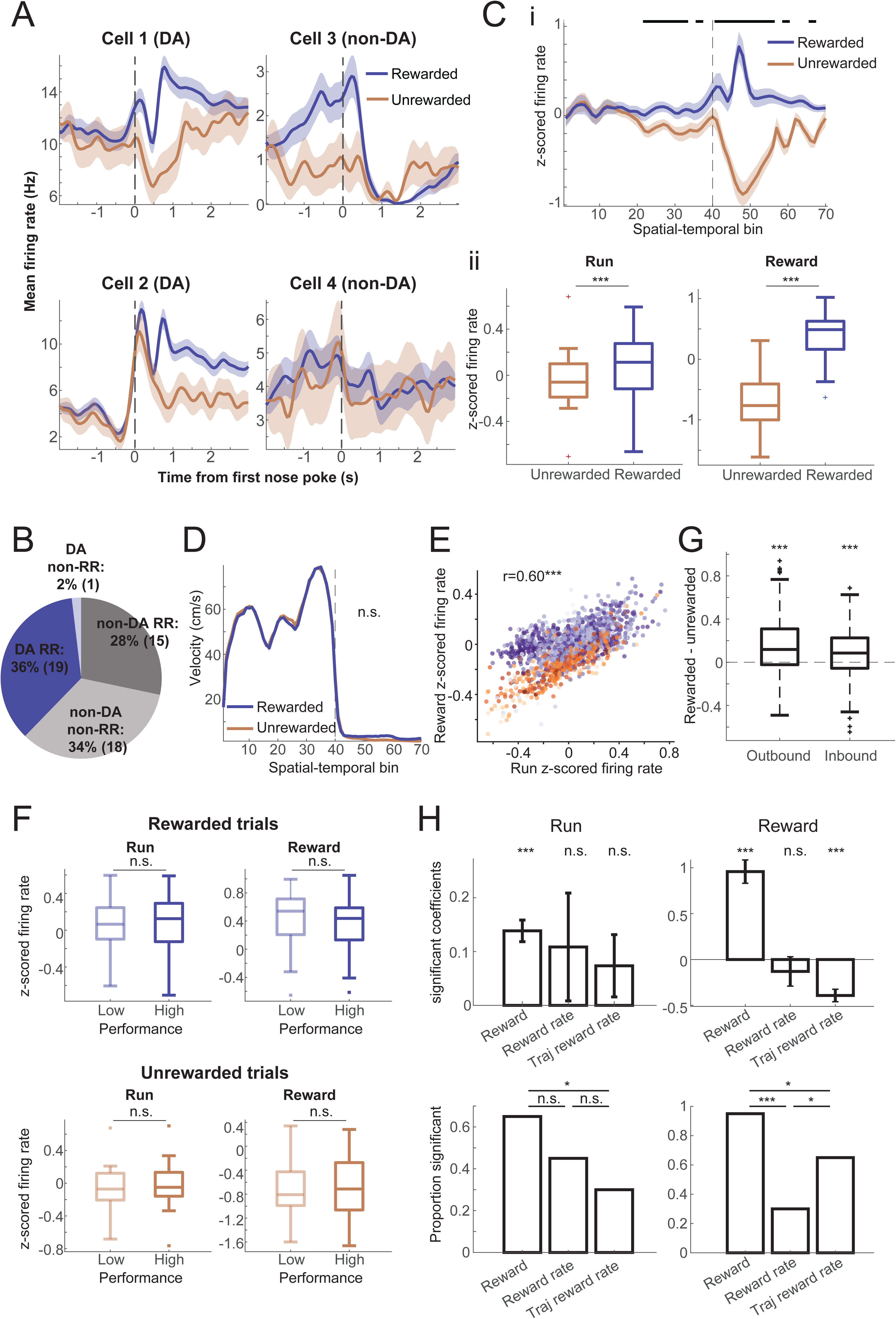
DA spiking activity mirrors reward prediction error. A. Illustrative examples showing that DA neuronal firing activity at reward wells signals robust differences between rewarded and unrewarded outcomes. Left: example photo-tagged TH+ neurons; right: example non-DA neurons. Line: trial-averaged firing rate; shaded area: standard error. B. Percentage of cell types and reward responses in VTA. RR: reward responsive at 200-1200ms after first nose pokes. DA RR 36% (19/53), DA non-RR 2% (1/53), non-DA RR 28% (15/53), non-DA non-RR 34% (18/53). C. (i) Population response of putative DA neurons to reward outcomes during reward approach and at reward well. Bar on top indicates spatial/temporal bins with p<0.05 after Bonferroni correction using paired rank tests. Vertical black dotted line is first nose poke/ reward at the reward well, also corresponding to transition from spatial to temporal bins. Spatial bins during approach to reward well are from 1-40 (5-6 cm per bin, 200-240 cm in total), and temporal bins after arrival at reward well are from 41-70 (100 ms per bin, 3 seconds in total). (ii) average z-scored firing rates of all putative DA cells during rewarded vs unrewarded trials. Left: during run, starting from leaving the last reward/nose poke to the beginning of the current trial’s reward/nose poke, rewarded median=0.11, unrewarded median = −0.060, p = 1.8×10^−4^; right: reward time (200ms-1200ms after the first nose poke), rewarded median=0.49, unrewarded median = −0.76, p = 5.6×10^−5^. Wilcoxon signed-rank test, n=20 cells. D. Control for running speed during trial. No significant differences are seen in running speed between rewarded and unrewarded trials (p = 0.12). E. Strong correlation of DA population firing rates during reward approach running and at reward outcome (r = 0.60, p = 6.8×10^−148^). Colors from light to dark indicate performance from low to high. Red: error trials; blue: correct trials. F. DA firing rates show no difference between low- and high-performance period. Top: rewarded trials (median z-scored firing rate during running: low performance 0.065, high performance 0.13, p=0.23; during reward outcome: low performance 0.54, high performance 0.44, p=1.00); bottom: unrewarded trials. (run: low performance −0.072, high performance −0.049, p=0.23; outcome: low performance −0.81, high performance - 0.71, p=0.65, Wilcoxon paired rank test, n=20 cells). G. Difference in DA firing rates for rewarded vs unrewarded trials during running is present for both inbound trials (median difference 0.085, p = 1.8×10^−6^, n = 178 trials) and outbound trials (median difference 0.12, p = 1.0×10^−13^, n = 217 trials). H. Multiple regression for DA neuron firing rates during run or reward outcome using the following parameters: current-trial reward, reward rate of the last 5 trials, reward rate the same trajectory for the last 5 visits. *Top*: mean significant coefficients. Current-trial reward: run: β = 0.14±0.02 (mean±sem), p=2.4×10^−4^; outcome: β = 0.95±0.13, p=1.6×10^− 4^; reward rate: run: β = 0.11±0.10, p=0.36; outcome: β = −0.13±0.16, p=0.56; same-trajectory reward rate: run: β = 0.074±0.058, p=0.16; outcome: β = −0.39±0.07, p=2.4×10^−4^, Wilcoxon signed rank test. *Bottom*: Proportions of DA neurons modulated by each factor. Run: current reward 65% vs reward rate 45%: χ^2^=1.6, p=0.20; current reward 65% vs same-trajectory reward rate 30%: χ^2^=4.9, p=0.027; reward rate 45% vs same-trajectory reward rate 30%: χ^2^=0.96, p=0.33. Outcome: current reward 95% vs reward rate 30%: χ^2^=18.0, p=2.2×10^−5^; current reward 95% vs same-trajectory reward rate 65%: χ^2^=5.6, p=0.018; reward rate 30% vs same-trajectory reward rate 65%: χ^2^=4.9, p=0.027. Fisher’s exact test, n=20 cells.

The influence of behavioral factors on each DA cell’s single-trial firing rates was estimated using multiple linear regression.

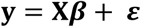

**y** is the mean firing rate of each trial during either running or the first 3 seconds of the outcome period. **X** is the estimated behavioral factors for each trial, which included: reward, binary outcome, rewarded or unrewarded; reward rate, an estimate of mean reward rate using the last 5 trials’ outcome; trajectory reward rate, an estimated reward rate of a certain trajectory, using outcomes of the last 5 times that the animal took the same trajectory. This process is repeated for all DA neurons (n=20), for both the running and reward outcome firing rates. The resulting significant coefficient values and the proportion of significant coefficients were summarized in **Figure 4H**.

## Results

We recorded neuronal activity simultaneously from the dCA1, PFC (anterior cingulate cortex and prelimbic regions) and VTA in adult male Th-Cre rats (n=4) using tetrodes in PFC and HPC and optical fibers surrounded by tetrodes in VTA (**Figure 1E-G**, see **Materials and Methods**), while animals performed a non-cued spatial rule switching task (**Figure 1A**). Both rules in the switching task consist of two types of trajectories with different memory demands: inbound trajectories that are rewarding at the home location regardless of the animals’ travel history; and outbound trajectories that only reward animals when they choose the different arm from last non-home visit, requiring working memory. For rule 1 (standard W/-M maze alternation task), the middle arm is the home location and animals alternate between left and right arms for outbound trajectories. For rule 2, the home location is switched to the left arm, and the middle and right arms become outbound destinations. Animals learned the two rules and were subsequently trained to switch between the two rules with solely the feedback of reward outcomes, and no external cues signaling the rule switch (**Figure 1B**). Animals were first trained on Rule 1 and subsequently on Rule 2. After 7-10 days of learning and training, animals could switch between the rules 3-4 times within a recording day. Rule switching was triggered manually during a session after a threshold performance (>80% correct) was achieved on the current rule. Upon unexpected trial outcomes, animals adapted their behavioral choices quickly to obtain more rewards, evident in decreasing numbers of perseverative errors over time (**Figure 1D**). It took 17 trials on average to reflect a Rule 2 to 1 switch in behavior, and 22 trials for Rule 1 to 2 (**Figure 1C**). The Rule 1 to 2 direction was typically more difficult for animals to achieve due to spatial asymmetry of rule 2. During each day of recording, animals ran the tasks in 3-4 20-30 min sessions, which were interleaved by 20-40 min sleep sessions. In the first or last sleep session, opto-tagging was performed to identify putative DA neurons in VTA (**Figure 1H**). An example raster during one trial is shown in **Figure 1I**. Position and speed plots show animal motion and immobility at the destination reward well, and raster plots show corresponding patterns in the three regions. Note that CA1 and PFC firing activity spans the spatial locations on the track. A total of 143 CA1 neurons, 128 PFC neurons and 53 VTA neurons (20 TH+ neurons) from 4 animals, one recording day for each, were included in this study (**Table 1**).

**Table 1:**
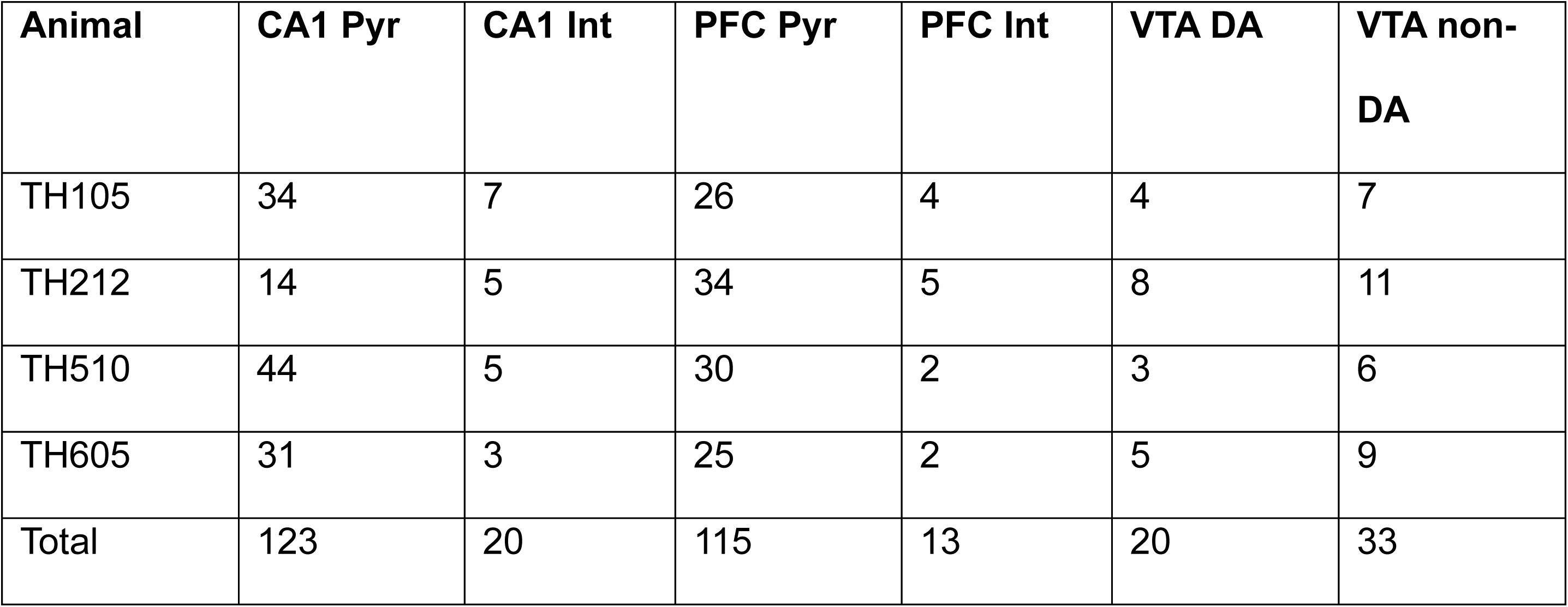
Clustered cell counts by brain region and cell type.

| Animal | CA1 Pyr | CA1 Int | PFC Pyr | PFC Int | VTA DA | VTA non-DA |
| --- | --- | --- | --- | --- | --- | --- |
| TH105 | 34 | 7 | 26 | 4 | 4 | 7 |
| TH212 | 14 | 5 | 34 | 5 | 8 | 11 |
| TH510 | 44 | 5 | 30 | 2 | 3 | 6 |
| TH605 | 31 | 3 | 25 | 2 | 5 | 9 |
| Total | 123 | 20 | 115 | 13 | 20 | 33 |

### CA1 and PFC neurons show differentiated activity across rules

CA1 and PFC neurons show spatially modulated firing activity (O’Keefe and Dostrovsky, 1971; O’Keefe, 1978; Hyman et al., 2010; Zielinski et al., 2019), and their features have also been implicated in rule and context representations (Wood et al., 2000; Eschenko and Mizumori, 2007; Griffin et al., 2007; Rich and Shapiro, 2009; Durstewitz et al., 2010; Ferbinteanu et al., 2011; Karlsson et al., 2012; Powell and Redish, 2016; Guise and Shapiro, 2017; Hasz and Redish, 2020). We therefore first asked the question if the firing activity of CA1 and PFC neurons can reflect the current rule by showing rule-specific activity on trajectories. As firing activity of VTA neurons showed much weaker spatial modulation (**Figure 2E**), we focused only on CA1 and PFC for this analysis. Single-unit remapping was investigated using firing activity on the two common trajectories that lead to reward for correct trials for both rules (*Left-to-Center*: inbound for Rule 1 and outbound for Rule 2, and *Center-to-Left*: outbound for Rule 1 and inbound for Rule 2), as lack of reward is known to destabilize firing activity on mazes (Krishnan et al., 2022). We found cells showing remapped activity across the two rules in both CA1 and PFC regions (**Figure 2A**; rules are run in alternating blocks with two blocks for each rule). Some cells show rate remapping (**Figure 2A**, first example on left, CA1 excitatory cell), and some cells exhibited relocation of spatial firing to a different part of the track (CA1 and PFC excitatory cell examples). Additionally, reward-associated firing after the same physical trajectory but in different rule contexts also showed robust differences (examples with significant differences in **Figure 2B**). These firing changes are not a result of recording drift over time, as similar patterns were observed both earlier and later in the day, with interleaved rule blocks. Further, trial-wise cell firing vectors also exhibited increases in correlation with stable-performance firing vectors of the current rule as behavioral performance improved, and become less similar (lower correlation) after the rule changed (**Figure 2C**). Overall, we found that 27.3% of hippocampal CA1 cells and 25.8% of PFC cells show significantly differentiated activity during running based on the current underlying rule, and 14.7% CA1 and 14.8% PFC neurons during reward. There was no significant difference in the ratio of rule-modulated neurons across regions (**Figure 2D**).

### CA1 and PFC ensemble representations distinguish the rules and show transitions during rule switching

We next investigated ensemble activity changes across rules in CA1 and PFC, and the dynamics of these changes in relationship to behavior. Trial-by-trial population vector similarity was computed and aligned to the first trial after rule switching, shown in the correlation plots in **Figure 3A**. Similar to previous research findings (Hasz and Redish, 2020), the firing pattern of both regions stabilized toward the end of running one rule (seen as increase in correlation with neighboring trials within a rule block) and quickly destabilized after changing to the other rule (**Figure 3A**). We adopted a newly developed dimensionality-reduction method CEBRA (Schneider et al., 2023), utilizing contrastive learning to visualize the difference of high-dimensional neural data across rules. For a common trajectory across the two rules (*Left-to-Center*), PFC 3D embeddings using CEBRA qualitatively showed consistent separation across rules for all animals during stable performance (visualized in **Figure 3B (ii)**, top row). These embeddings used training and test datasets within animals. We also used multi-session training that sampled training data from all animals followed by testing within animals. The embeddings from this method revealed a similar structure across animals, implying animal-invariant features in the PFC ensemble data for rule representations (**Figure 3B (ii)**, bottom row). CA1 single-session embeddings showed similar separations between rules compared to PFC. However, much weaker separations were observed in multi-animal embeddings, suggesting a more unique latent space of CA1 rule-specific activity for individual animals (**Figure 3B (i)**). As animals switched from stable performance of one rule to another, we also observed a systematic transition in the manifold space (example PFC manifold transitions over trials in **Figure 3C**).

In order to quantify these changes, a KNN decoder was trained to decode the underlying rule using single-animal embeddings for each brain region (see **Materials and Methods**). Rule decoding probabilities using CA1 and PFC data correlated highly with animals’ behavior strategy (**Figure 3D, E**). Upon further inspection, behavior strategy showed stronger correlation with PFC rule decoding than with CA1 decoding (**Figure 3E**). The decoding probability of the current rule increased with behavioral performance for both PFC and CA1 (**Figure 3F**, mean decoding accuracy for low performance: CA1 0.38, PFC 0.41; mid performance: CA1 0.66, PFC 0.71; high performance: CA1 0.97, PFC 0.96; performance thresholds are reported in **Materials and Methods**). However, when training the decoder using multi-animal embeddings, CA1 ensembles performed much worse than PFC, again implying more consistent rule coding across animals in PFC and its stronger relevance to behavior than CA1 (**Figure 3F**).

We further examined the timing between neural representation change and behavior strategy shift upon rule switch. The behavior and neural changes align with each other, with neural representations in both CA1 and PFC transitioned more often before behavior strategy than after (**Figure 3H, I**), with no timing difference seen between CA1 and PFC ensembles. On average, it took more trials to switch from rule 1 to rule 2, both in the neural representation and behaviorally (**Figure 3G**), potentially due to asymmetric working memory components in rule 2.

### VTA DA firing activity reflects and predicts reward outcomes

As midbrain DA plays a crucial role in reward signaling, we aimed to understand VTA DA spiking activity features during rapid rule switches guided by unexpected reward outcomes. We used an opto-tagging strategy in TH-Cre rats, which has been utilized in previous studies (Witten et al., 2011). Opto-tagging was performed during the first or last sleep session for each day of recording to not interfere with activity during task running. We recorded a total of 53 VTA neurons, out of which 20 photo-tagged cells were identified as putative DA neurons (with firing rates, mean ± std, 7.7 ± 3.0 Hz). Consistent with a vast body of literature, putative DA neurons showed increased firing rates when rewards became available, temporally aligned to first poke at reward well, and decreased firing rates during reward omission (**Figure 4A**, median z-scored firing rate for rewarded trials: 0.49, unrewarded trials: −0.76, p = 5.6×10^−5^ Wilcoxon signed rank test). Non-tagged neurons exhibited a diverse range of activity, including 45.5% (15/33) of neurons responding to reward outcomes (**Figure 4B**). Further, we also observed that DA neurons had higher firing rates during running for trials that lead to rewards in comparison to unrewarded trials (**Figure 4C**, median z-scored firing rate for rewarded trials: 0.11, unrewarded trials: −0.060, p = 1.8×10^−4^). This difference became apparent and significant halfway through running on the trajectory, just after the choice point where the decision was made, similar to previous findings of DA release in striatum, and is likely related to reward expectancy/ uncertainty (Howe et al., 2013). The running speed profile on the track was similar between rewarded and unrewarded trials, suggesting the DA firing rate difference is not caused by a change in motivation or vigor (**Figure 4D**).

Based on the proposed link between DA spiking and reward prediction error, it is predicted that for rewarded trials, as behavioral performance improves, DA firing rates at the reward will decrease and DA firing rates during run will increase; and such trends would be the opposite for unrewarded trials (Watabe-Uchida et al., 2017). However, we observed a strong positive correlation between run and reward firing rates (r = 0.60, p = 6.8×10^−148^), and no obvious relationship between firing rates of either state to performance (**Figure 4E**). We further tested this by separating the performance by trials into low and high performance halves, and found no difference in the firing rates across performance stages for either rewarded or unrewarded trials (n=20 cells, median z-scored firing rate for rewarded trials: run: low performance 0.065, high performance 0.13, p=0.23; outcome: low performance 0.54, high performance 0.44, p=1.00; unrewarded trials: run: low performance −0.072, high performance −0.049, p=0.23; outcome: low performance −0.81, high performance −0.71, p=0.65, Wilcoxon paired rank test, **Figure 4F**).

In addition, we wanted to investigate if the DA reward prediction signal is present for working memory error as compared to rule/perseverative error that occurs after a rule switch. At each time of rule switching, two trajectories became unrewarding under all conditions and two new ones became potentially rewarding when the sequence of arm visits was correct. For example, when rule 1 changes to rule 2, trajectories between the center and the right arms are no longer rewarding. Instead, left-to-right and right-to-left ones yield rewards. We classified the trials that animals ran on the newly unrewarding trajectories as perseverative errors, whereas the trials that animals made a wrong outbound choice for the current rule as working memory errors. If the DA firing activity only reflects a model-free system tracking overall reward probability, we would expect a difference only between perseverative errors and correct trials, but not for individual decisions dependent on working memory. We found that the prediction for reward outcome was present in DA firing regardless of the error type, even when comparing error trials to nearest correct trials of the same outbound/inbound trajectory type (**Figure 4G**, median z-scored firing rate difference between rewarded and unrewarded trials: outbound 0.12, p = 1.0×10^−13^, n = 217 trials; inbound 0.085, p = 1.8×10^−6^, n = 178 trials).

To summarize the influence of reward conditions and histories on DA firing rates, we fitted multi-regression models for each cell’s spiking rates during running and outcome periods, separately. The most significant factor was found to be the current trial’s reward outcome (running: mean±sem coefficients: 0.14±0.02, significant for 65% of DA neurons; outcome: 0.95±0.13, 95%), in comparison to a weaker impact of reward history (running: 0.11±0.10, 45%; outcome: −0.13±0.16, 30%) or same-trajectory reward history (running: 0.074±0.058, 30%; outcome: −0.39±0.07, 65%), for both running and reward states (**Figure 4H**). Note that current trial reward outcome is a significant contributor to the approach run firing rate, and not just for reward well firing rate that is expected.

### DA reward predicting feature develops after rule switching and coordinates with PFC rule representation transition before behavior strategy adaptation

Our findings in PFC/CA1 rule coding and DA reward signals led us to the question -- if the firing rates of DA neurons predictive of reward outcomes develop as animals gather evidence of a rule change. Firing rates upon reward delivery were consistently higher than at reward omission (**Figure 5A (iv)**); however, the difference in DA neuron firing between rewarded and unrewarded trials during running only developed a few trials after the rule switch was implemented (**Figure 5A (v)**), and a few trials prior to behavioral strategy switch (**Figure 5A (i)**). We further investigated whether this DA firing activity is coordinated with rule representation changes in CA1 and PFC. When aligned to the trial where PFC started to decode the current rule, DA firing rate difference during run between rewarded and unrewarded trials ramped up robustly toward this change point (**Figure 5B**, bottom). The ramp up in DA firing rate difference was qualitatively more robust and consistent for PFC representation switch rather than when aligned to CA1 representation switch (**Figure 5C**, bottom). Animals’ behavior strategies adapted accordingly shortly after (on average 4 trials) the observed PFC rule representation switch and DA prediction emergence (**Figure 5B**, top). In addition, the DA firing activity difference between rewarded and unrewarded trials during maze running did not stay high as animals reached good, stable performance for the new rule. We observed a positive correlation between the DA firing rate difference and the rate of PFC decoding probability change, which may suggest a gating mechanism of value/belief update (**Figure 5D**). In summary, the reward predictive/expectancy property of DA firing during running is acquired post rule switch and is coordinated with the changes of PFC rule representation and behavior strategy.

**Fig 5:**
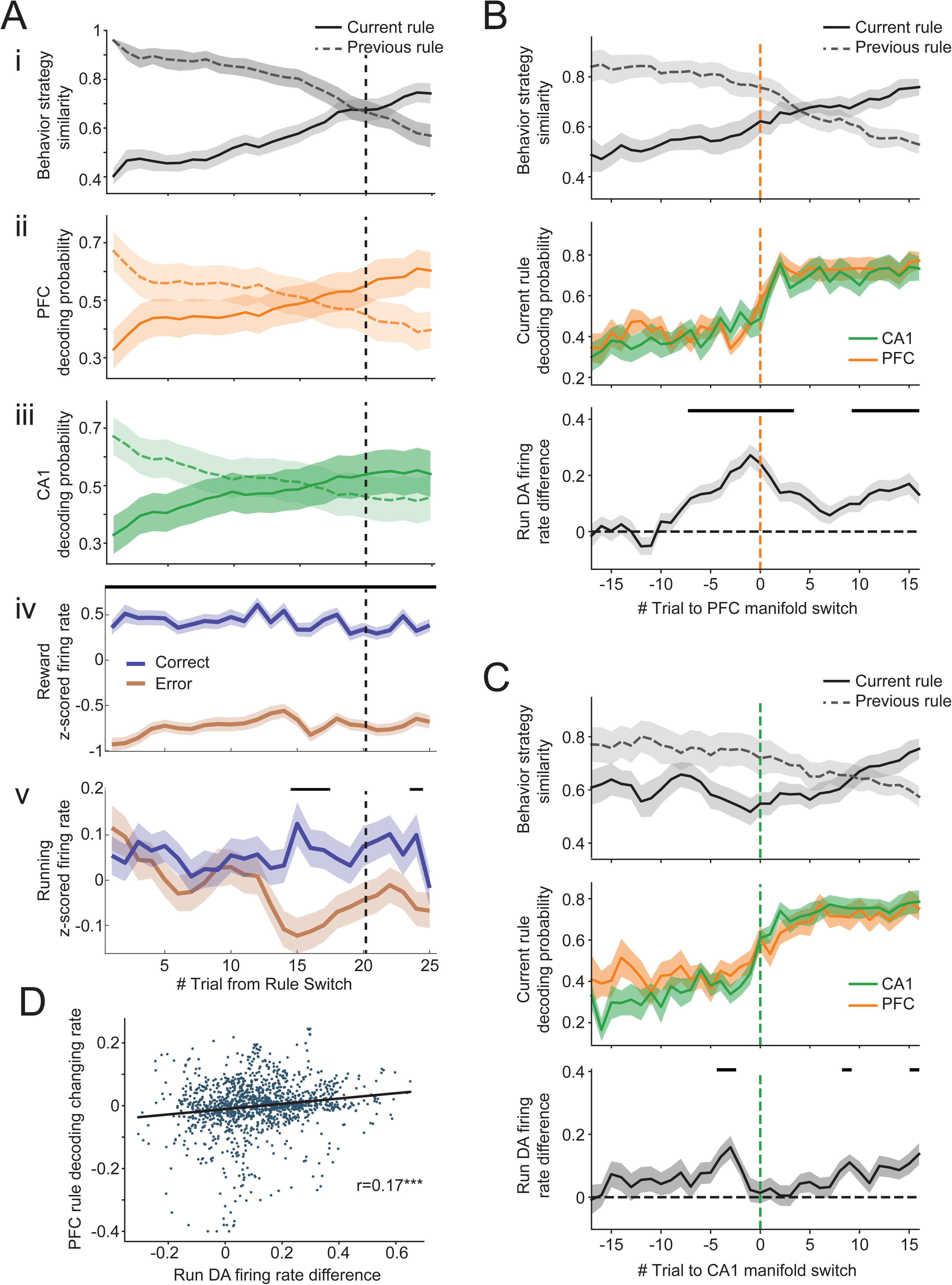
DA spiking activity relationship with neural manifold change and rule switching behavior. A. Behavior strategy (i), PFC (ii) and CA1 (iii) decoding probability changes aligned to rule switching trials. (iv-v). Difference in DA neuron firing rates for rewarded vs unrewarded trials at well locations (iv) and during running (v), plotted as a function of trials aligned to beginning of rule switch. Black bars indicate significant difference between rewarded and unrewarded trials with p<0.05 after Bonferroni correction. Vertical black dotted line across sub-panels denotes behavioral strategy switch trial. B. Behavior strategy, CA1/PFC decoding probability changes and DA firing rate differences between rewarded and unrewarded trials, aligned to decoded rule representation switch in PFC (vertical orange dotted line). Top: Behavior strategy similarity to optimized behavior of current and previous rules. Animals’ behavior adapted to the new rule on average 4 (IQR: −3.75-10.5) trials after PFC showed correct rule decoding. Middle: CA1 and PFC decoding accuracy of current rule. Bottom: DA firing rate difference for rewarded vs unrewarded trials during running. Black bars above indicate trials showing significant difference with p<0.05 after Bonferroni correction. C. Same as B but aligned to CA1 rule representation switch (vertical green dotted line). D. DA firing activity difference between rewarded and unrewarded trials is correlated with changing rate of PFC manifold (Pearson correlation r = 0.16, p = 2.8×10^−9^, n=1361 trials).

## Discussion

In the current study, we examined rule-related firing properties in the hippocampus and PFC, VTA DA spiking activity during rule switching, and the temporal relationships between cognitive rule representation, reward prediction signal and behavior adaptation. The mechanisms of rule representation and reward prediction have been generally examined separately, but it is vital to gain an integrated understanding of dynamics and temporal coordination between these mechanisms for supporting cognitive flexibility and behavioral adaptation. During rule switching, the presence or absence of rewards after specific actions can serve as feedback for organisms to reevaluate their behavior policy. A conflict between belief and outcome leads to a switch in internal rule representations, which can guide correct behavioral actions for the changed context. We therefore reasoned that there must exist some sort of temporal coordination between reward prediction and rule representation dynamics over the course of a few trials after rule switching, which will determine the time course of behavioral adjustments, thus allowing animals to overcome perseveration and adapt to the new context with appropriate actions. We found that PFC and hippocampal CA1 population activity represented rule context. VTA DA firing activity responded to and predicted reward outcomes. The timings of the switches in PFC-CA1 rule representation and the emergence of correct reward prediction from VTA were coordinated. Further, these changes in PFC, CA1 and VTA DA firing activity led the behavioral strategy transition, suggesting coordinated network activity switches to support behavioral adaptation.

We implemented a spatial rule switching task in a W/M maze, probing behavioral flexibility and working memory at the same time in the exact same environment. Rule 1 comprises a traditional W-maze alternation rule with the center arm as home arm as in previous studies (Frank et al., 2000; Jadhav et al., 2012, 2016; Tang et al., 2017; Maharjan et al., 2018; Shin et al., 2019), and the home arm switched to the left arm for Rule 2, requiring the animals to change their strategy for optimal trajectory sequences across the rules. Notably, both rules incorporate a spatial working memory component (outbound component), requiring the animals to choose the opposite arm from previous choice when embarking from the home arm. The return-to-home-arm trajectory is the inbound component with a spatial reference memory demand (Kim and Frank, 2009). The nature of the task assigned different memory demands of overlapping trajectories in the two rules, which allowed us to compare animal behavior, CA1/PFC representations and DA signaling for reward prediction during rule switch.

We first examined the differences of rule representation between the hippocampal CA1 and PFC. Spiking activity changes associated with rule/context were observed in both brain regions (Guise and Shapiro, 2017; Hasz and Redish, 2020), with single neurons exhibiting remapping on the same trajectories across the two rule contexts. Examining ensemble coding using manifold analysis and decoding to current rule context revealed more robust and consistent coding of rule contexts across animals in PFC, suggesting that similar neural representations may develop in PFC to support two rule contexts in the same physical trajectory space on mazes. Rule representations in PFC appeared to shift over the course of a few to tens of trials after rule switching, prior to observation of a behavioral strategy switch.

We observed more consistency across animals in PFC ensemble decoding for rules, whereas CA1 ensemble representations were more animal-specific (**Fig. 3B**). In terms of timing, we did not see a significant difference between PFC and CA1 in terms of neural transition times as compared with behavioral transition times, in terms of number of trials, although there was some variability across regions (**Fig. 3H, I**). However, the DA prediction differences that emerged after rule changes were more robustly aligned to PFC transition times than to CA1 (**Fig. 5B-C**), and PFC transition times were more closely aligned to behavioral strategy switch (**Fig. 3E**). Therefore, although there were no significant timing differences between PFC and CA1, this suggests more consistency in PFC neural transitions and behavioral transitions.

We observed that VTA DA neurons participated in reward prediction and this rule-specific feature developed quickly after rule switching. Previous literature has often focused on model-free RL reward prediction error; here, we instead tried to understand how midbrain DA neuronal firing activity changes during a rule switching task that requires working memory. DA neuronal firing rates were highly correlated between running and reward outcome in a trial, which somewhat resembles previous findings in (Coddington and Dudman, 2018). In addition, we did not find a correlation between DA spiking activity and performance at the population level. These results contrast with the prediction based on classical conditioning that firing activity increases toward cue/beginning of a trial and slowly diminishes upon reward delivery (Schultz et al., 1997; Fiorillo et al., 2003; Amo et al., 2022). Rather, our findings suggest that DA signaling may incorporate rule-based and working-memory dependent prediction information, potentially from cortical input.

We found a delay in animals’ behavioral strategy adjustment in comparison to rule representation and reward prediction. Animals continued to sample from wrong trajectories, although at a lower frequency, after manifolds shifted to the new rule and DA signal could predict the reward outcome. This can be explained by simply reserving a small chance for random exploration, or perhaps by the theory of active inference (Friston et al., 2012, 2014). In this theory, the dopamine signal minimizes surprise instead of cost. Therefore, sampling from extra trials could help confirm animals’ beliefs if correct and eventually stabilize behavioral sequences.

How is information processed in these regions? Does VTA receive reward, expectation or reward prediction error (RPE) information from other regions? Lisman and Grace (2005) suggested that novelty information is conveyed to VTA from the hippocampal formation through an indirect route, possibly with inhibitory afferents from accumbens and ventral pallidum. In return, DA fibers innervate dorsal hippocampus and thus enhance LTP (Huang and Kandel, 1995; Otmakhova and Lisman, 1996; Li et al., 2003; Rosen et al., 2015; Sayegh et al., 2024). In terms of DA-cortical interactions, VTA receives abundant prefrontal glutamatergic projections (Geisler et al., 2007), and their impact on reward signaling is multifaceted. Input from frontal cortices is critical for conveying predictive and incentive features of a cue (Pan et al., 2021). PFC conveys belief state information to the dopaminergic system and affects RPE computation when hidden states are involved (Babayan et al., 2018; Starkweather et al., 2018). Additionally, Amo et al. (2024) showed glutamatergic input to VTA already carried RPE signal, challenging the classic view of local computation of RPE in VTA by combining different aspects of reward and expectation from glutamatergic and GABAergic inputs (Kawato and Samejima, 2007; Keiflin and Janak, 2015). An intriguing future direction of research could focus on dissecting computational features or components of RPE in these brain regions involved in the reward circuit, particularly when multiple cognitive demands are present.

It is important to differentiate between DA firing activity and DA release in downstream area. We did not observe an obvious increase in DA spiking activity during goal approach. However, downstream release in striatum has been reported to ramp up towards goal (Howe et al., 2013; Hamid et al., 2015), suggesting partially dissociated activity potentially caused by local modulation (Mohebi et al., 2019). It would be interesting to examine if the working-memory dependent reward prediction signal is present in the DA release profile as in VTA DA spiking activity, and if so, how that influences its downstream striatal and cortical areas.

The exact mechanism of information transfer and cooperation requires further investigation. For example, Fujisawa and Buzsáki (2011) proposed a 4 Hz rhythm orchestrating neuronal activity in the hippocampus, PFC and VTA during a similar working memory task. DA administration in PFC increased HPC-PFC theta coherence and PFC phase locking to CA1 theta (Benchenane et al., 2010). PFC theta sequences selectively coordinate with CA1 theta sequences depicting future trajectory choices (Tang et al., 2021). Therefore, altered DA neuron spiking activity might be impacted by the online processes of action evaluation and selection during running. Another possible mechanism is through coordinated reactivation/replay. It has been shown that hippocampal reactivation is associated with increased firing activity of VTA cells as a potential mechanism for memory consolidation (Gomperts et al., 2015). CA1-PFC coordinated replay is biased toward behavioral choice and therefore can potentially participate in working memory and decision making (Shin et al., 2019). This hippocampo-cortical/subcortical temporal coordination during offline states may serve as substrates for value updates, memory consolidation, action evaluation and planning. Our study focused on the behavioral time scale, but investigating simultaneous activity on a shorter time scale (50-200 ms) can potentially uncover detailed mechanisms of inter-regional communication.

## Conclusions

To understand the temporal coordination between cognitive systems and reward systems, we recorded from the rat hippocampal CA1, PFC and VTA during a rapid rule switching task. CA1 and PFC exhibited rule representation in their population activity. DA neuron firing activity in VTA gradually became predictive of reward outcomes after rule switching, and this predictive feature developed together with the PFC representation transition to reflect the new rule. These neural representations changed in advance of behavioral adaptation by a few trials. Together, our study revealed a synchronized update of information across PFC, HPC and VTA that potentially supports behavioral flexibility.

## Acknowledgements

This work was supported by the National Institutes of Health (R01MH112661) to S.P.J. and a T90 training grant (R90DA033463). We would like to thank Dr. Naoshige Uchida, Dr. Joshua Berke and Dr. Ali Mohebi for the protocol of the opto-tagging experiment, Dr. John Bladon and Audrey Hooker for piloting the W-track rule switching task, Dr. Justin Shin in assisting in implant surgeries, Dr. Wenbo Tang, Dr. Jacob Olson, Dr. Blake Porter and other lab members in providing feedback on the research project.

